# Presynaptic GABA_B_ autoreceptors suppress neurotransmitter release during repetitive stimulation via the Gβγ-SNARE pathway

**DOI:** 10.64898/2025.12.10.693514

**Authors:** Zack Zurawski, Iris Lu, Mariana Potcoava, Claire E. Delbove, Christian J. Peters, Heidi Hamm, Simon Alford

**Affiliations:** Department of Anatomy and Cell Biology, University of Illinois at Chicago, Chicago, IL; Department of Pharmacology, Vanderbilt University, Nashville, TN

## Abstract

GABAergic signaling provides the brain’s primary inhibitory mechanism with defects linked to epilepsy, anxiety, depression, insomnia, schizophrenia and neurodegeneration. A key regulatory mechanism is autoinhibition of GABA release during repetitive activity via presynaptic G_i/o_-coupled GABA_B_ receptors, supporting synaptic tuning and memory formation, and limiting neurotransmitter spillover. Exogenous GABA_B_ receptor agonists reduce presynaptic Ca^2+^ entry by inhibiting calcium channels. However, using transgenic mice expressing a mutant SNAP25 with diminished ability to bind Gβγ (SNAP25Δ3), we show that suppression by GABA_B_ autoreceptors requires intact Gβγ-SNARE interactions. Imaging of presynaptic Ca^2+^ transients in GABAergic axons showed no GABA-mediated autoreceptor suppression of Ca^2+^ entry during stimulus trains. In contrast, application of the exogenous GABA_B_ receptor agonist baclofen profoundly inhibited Ca^2+^ entry, which could be partially reversed by exogenously elevating cAMP, indicating a complementary role of inhibition of adenylyl cyclase. Baclofen reduced spontaneous IPSC frequency and amplitude and both effects were diminished in SNAP25Δ3 mice, consistent with inhibition at Ca^2+^ channels and SNARE complexes. Physiological GABA-mediated and exogenous GABA_B_ receptor activation thus produce distinct outcomes on GABAergic neurotransmission, indicating that synthetic drug application to neurons does not faithfully recapitulate endogenous signaling pathways. We conclude that endogenous rapid GABA_B_ autoreceptor signaling inhibits neurotransmitter release primarily by Gβγ-mediated inhibitions of SNARE mechanisms, whereas prolonged agonist application additionally suppresses Ca^2+^ influx via cAMP signaling.

## INTRODUCTION

Neuronal autoreception is a signaling mechanism by which a receptor at presynaptic or prejunctional nerve terminals is activated by neurotransmitter release from that same axon. This can allow for dynamic modulation of synaptic transmission during periods of high frequency neurotransmitter release. Many of these autoreceptors are inhibitory Gα_i/o_-coupled GPCRs^1^ that provide critical negative feedback by attenuating subsequent neurotransmitter release after an initial release event at that axon^2^. Specific autoreceptors have been proposed or identified for many types of neurotransmitters and neuromodulators, such as mGluR7 for glutamatergic neurons, α2a for adrenergic neurons, and 5-HT_1_ for serotonergic neurons acting both centrally and peripherally^3–5^. Their importance is apparent in all of these systems. For example, glutamate release in excitatory neurotransmission depolarizes the postsynaptic membrane via ionotropic glutamate (AMPAR and NMDARs). This is necessary for signaling and plasticity, but excessive release leads to cell death^6–8^. Metabotropic glutamate receptors attenuate this release.

By a similar mechanism, GABAergic inhibitory neurons utilize GABA_B_ receptors as autoreceptors on their presynaptic terminals to attenuate GABA release events during trains of action potentials^9,10^ . In hippocampal CA1 pyramidal cells, this shapes postsynaptic GABA-mediated responses during high frequency activity; GABA_B_ receptor-mediated inhibition of GABA release relieves a brake on excitation, enabling the induction of long-term potentiation (LTP)^11,12^ via enhanced activation of postsynaptic NMDA receptors. GABA_B_ autoreceptors can also act pathologically to silence inhibitory neurons without concomitant activation of GABA_B_ heteroreceptors on pyramidal cells, which can result in disinhibition and pathological seizure activity in the hippocampus^13–15^. Critically for cortical and hippocampal function, GABA_B_ autoreceptors fine-tune network excitability to keep competing excitatory and inhibitory signals in balance within time frames of milliseconds to seconds^16–18^.

As Gi/o-coupled GPCRs, GABA_B_ receptors are canonically thought to signal via inhibition of adenylyl cyclase^19^, causing activation of GIRK channels to produce hyperpolarization^20–23^ or reduction of Ca^2+^ fluxes through voltage-gated Ca_V_2 channels^24–27^ , but regulation downstream of Ca^2+^ channel also occurs.. As a heteroreceptor, GABA_B_ has been shown to act downstream of Ca^2+^ entry via the Gβγ-SNARE pathway at glutamatergic synapses onto medium spiny neurons^28^, but not at other glutamatergic synapses^1,24^. The extent to which GABA_B_ autoreceptor signaling occurs at or downstream of voltage-gated Ca^2+^ channels at inhibitory synapses onto pyramidal synapses is currently unknown. Using fast intracellular genetically encoded calcium indicators^29^ and genetically encoded synaptic biosensors specific for GABA^30^during high resolution live cell imaging^31^, it is now possible to distinguish between these mechanisms at the level of individual synapses within acute *ex vivo* tissue. We show that exogenous agonist used to probe presynaptic effects of GABAB receptors markedly inhibit GABA release by inhibiting presynaptic Ca^2+^ entry, but that endogenously released GABA acts via a distinct effect of Gβγ directly at the SNARE complex.

## RESULTS

### GABA_B_ receptors autoinhibit GABA release

To quantify autoreception, transsynaptic GABA signaling is manipulated experimentally by engaging presynaptic mechanisms. To validate this, we compared hippocampal IPSCs evoked by repetitive stimulation of presynaptic GABAergic terminals with events evoked after application of the exogenous agonist baclofen. These were recorded in acute brain slices using patch clamp of hippocampal CA1 pyramidal neurons, while either stimulating stratum radiatum or infusing baclofen into bath solutions (Figure 1A). CA1 pyramidal neurons were whole cell voltage clamped and held at -50 mV. The whole cell patch solution contained Cl^-^ as the principal anion (129 mM), causing GABA_A_-mediated postsynaptic events to exhibit inward currents. GABAergic inhibitory postsynaptic currents (IPSCs) were evoked by local extracellular stimulation within 200 µm of the recorded neuron (Figure 1A). Repetitive stimuli were applied in trains of 5 pulses from 2 Hz to 20 Hz and recordings were made in 6-cyano-7-nitroquinoxaline-2,3-dione (CNQX; 5 µM) and 2-amino-5-pentanoic acid (AP5; 50 µM) to block glutamatergic transmission and isolate monosynaptic IPSCs. Recordings revealed a short-term synaptic depression in IPSC amplitude, as measured by reduction in peak amplitude between the first and subsequent IPSCs in the train (Figure 1B,D) as previously reported^10,32^.

**Figure 1:**
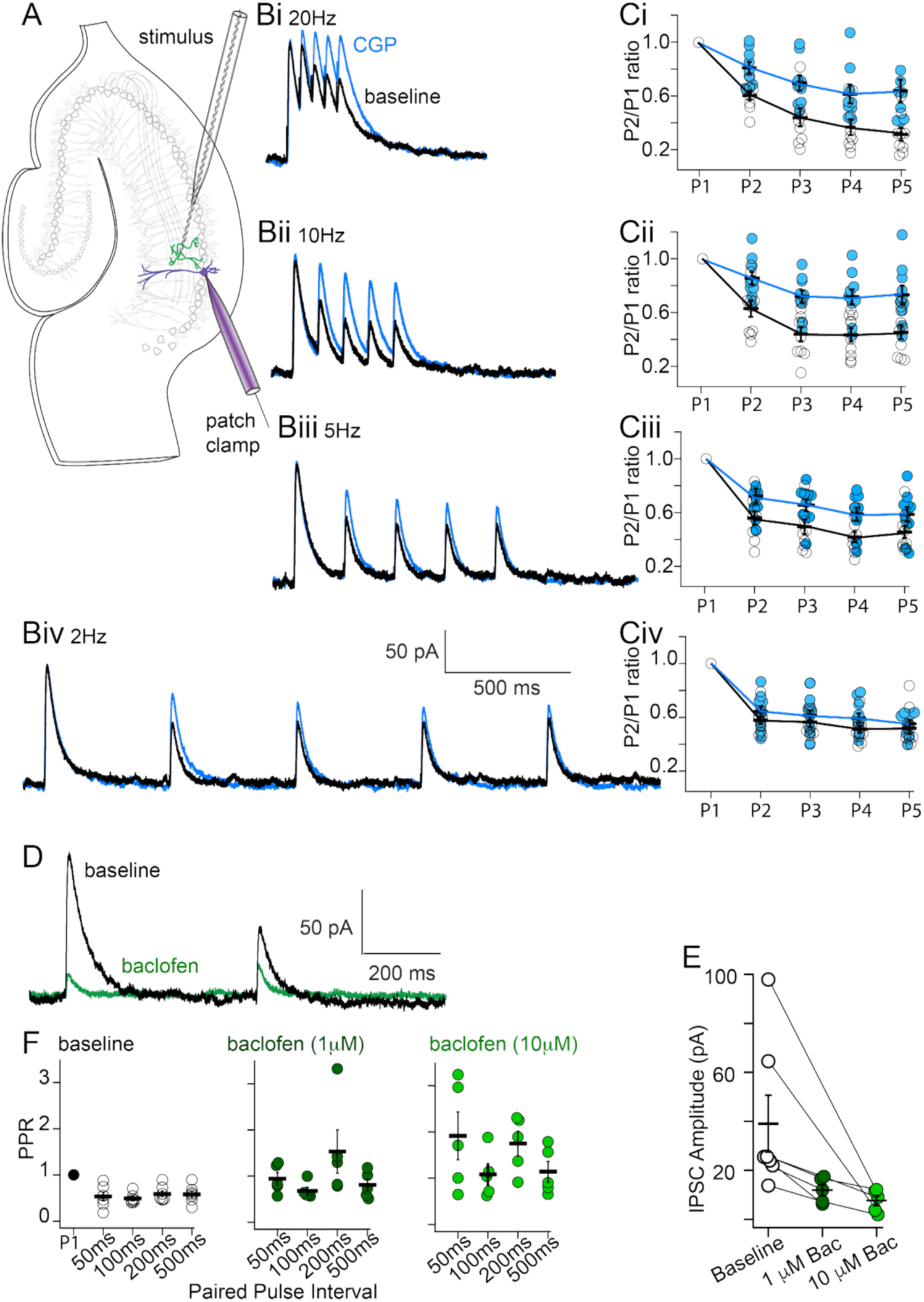
GABA_B_ activation by autoreceptors or the exogenous agonist baclofen inhibits GABAergic neurotransmission at CA1 pyramidal cells. A. Schematic diagram of electrophysiological assay for measuring electrically evoked isolated inhibitory postsynaptic currents, in which CA1 pyramidal neurons are held in the whole-cell voltage-clamp configuration in the presence of 50μM AP5 and 5μM CNQX. B. Representative traces of 5-shock IPSC prior to and in the presence of 10μM of the GABA_B_ antagonist CGP 55845 (in blue) at 20 Hz (Bi) 10 Hz (Bii) 5 Hz (Biii) or 2 Hz (Biv). C. Scatterplot of the relative ratio of peak amplitudes for each subsequent stimulus to the first at a frequency of 2 Hz. Connecting line plotted at arithmetic mean of each genotype. The experiment was performed 7 times for 6 biological replicates. The effect of CGP 55845 was significant for 20 (Ci) and 10 Hz (Cii), as tabulated via multiple comparisons tests with the Bonferroni correction after 2-way ANOVA with repeated measures. D. Representative traces of 2-shock IPSC train prior to and in the presence of the GABA_B_ agonist baclofen (10μM, in green) at 2 Hz (as in Biv) P-values obtained from two-tailed Student’s t-test. E. Scatterplot of amplitudes for the first IPSC in the train , treated with 1 or 10μM baclofen in the bath. The experiment was performed 7 times for 6 biological replicates. F. Paired-pulse ratios for the second IPSC in the train in the presence or absence of baclofen (10μM, in dark green, or 1μM, in light green), at paired pulse intervals of 50-500ms.

To verify that this IPSC suppression in repetitive pulse trains was due to GABA_B_ autoreceptor activation, we blocked GABA_B_ receptors by bath application of the GABA_B_ antagonist CGP 55845^33^. Application of 10 µM CGP 55845 did not alter the amplitudes of the first IPSC in the train (Supplemental Figure 1), but significantly diminished suppression of subsequent IPSCs in trains at 2, 5, 10 and 20 Hz (Figure 1C). These effects are consistent with short term depression of IPSC amplitude resulting from GABA_B_-mediated autoreceptor-mediated inhibition during repetitive stimulation^10,32^.

We therefore expected that the GABA_B_ agonist baclofen would mimic aspects of GABA_B_ autoreceptor-mediated inhibition. Application of baclofen (1 – 10 µM) dose dependently inhibited IPSC amplitudes (Figure 1D, E). It also dose-dependently reversed paired pulse inhibition, consistent with an occlusion of the GABA_B_ autoinhibition (Figure 1F). However, the maximal inhibition of the response by baclofen was significantly larger than the autoinhibition mediated by trains of stimuli. Maximal inhibition in baclofen (at 10 µM) was to 12.60 +/- 2.97% of baseline compared to a maximal inhibition to 49.24 +/- 5.81% following 5 shocks of train stimulation at 10 Hz. The effect of baclofen on the dynamics of the IPSC was also different from that seen during stimulus trains: baclofen significantly increased the rate of decay of the responses, but not the rate of activation (Supplemental Figure 2), whereas no effect of repetitive stimulation was seen on IPSC decay rates for any responses where the decay rate could be adequately fit with an exponential. From these findings, we concluded that baclofen can inhibit evoked IPSC amplitudes to a greater extent than endogenous GABA-driven inhibition at physiologically relevant frequencies.

### Differential impacts of baclofen and endogenous GABA release on presynaptic Ca^2+^ signaling of inhibitory neurons

Electrophysiological and imaging studies have generated considerable evidence that presynaptic GABA_B_ receptor activation leads to inhibition of Ca^2+^ entry, which in turn is believed to represent the mechanism by which presynaptic GABA_B_ receptors inhibit GABA release. This is thought to occur via the binding of free Gβγ subunits to voltage-gated Ca^2+^ channels (VGCCs) to reduce action potential evoked presynaptic Ca^2+^ fluxes^34^. Evidence from bulk loading of dye into presynaptic Schaffer Collateral Commissural (SCCP) axons indicated that baclofen inhibits presynaptic Ca^2+^ entry to excitatory terminals^25^ Furthermore, line scanning confocal microscopy of GABA_B_ heteroreceptor function at single excitatory CA1-subicular synapses in the hippocampus has demonstrated GABA_B_ receptor-dependent inhibition of Ca^2+^ entry at individual glutamatergic terminals^24^.

To image GABA_B_ autoreceptor function in intact GABAergic terminals, AAVs containing the genetically encoded fast Ca^2+^ sensor jGCaMP8f under the control of the inhibitory neuron-specific enhancer mDlx^35,36^ were stereotaxically injected into the hippocampi of adult mice. Acutely harvested hippocampal slices were then imaged using epifluorescence microscopy. jGCaMP8f fluorescence was observed in whole inhibitory neurons, including the soma, dendrites and axons (Figure 2A). Axo-somatic inhibitory synapses from these neurons were observed in the cell body layer of CA1, and these presynaptic structures were used for subsequent functional assays. Using similar stimulator placement as shown in Figure 1, stimulus trains (2 to 10 Hz) were applied to the stratum radiatum and jGCaMP8F fluorescence collected at 100 Hz from fluorescent axon terminals, while avoiding inclusion of soma or dendrites of the presynaptic neuron within the recorded region. To block polysynaptic activation and Ca^2+^ fluxes from ionotropic glutamate receptors from adjacent dendrites, 5µM CNQX and 50µM AP5 were present in bath solutions.

**Figure 2.**
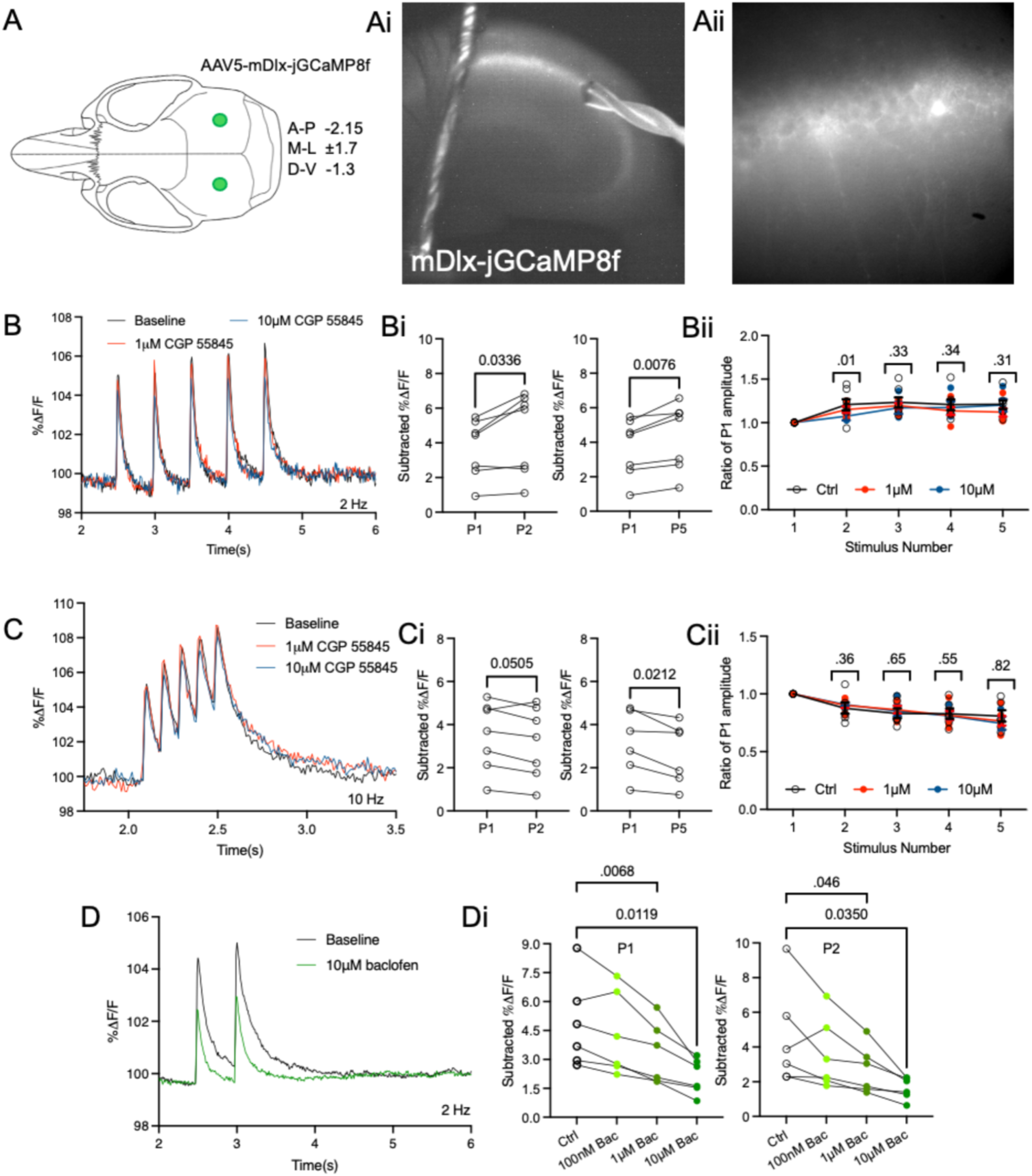
Baclofen, but not GABAB autoreceptor signaling, inhibits GABAergic neuronal Ca^2+^ fluxes. A. Visual schematic of stereotactic injection of AAV1-mDlx-jGCaMP8f-WPRE into mouse parenchyma. Ai. Acute mouse hippocampal slice expressing bright mDlx-jGCaMP8f fluorescence within and in close proximity to the stratum pyramidale imaged via epifluorescence microscopy, at 4x magnification. Retaining harp and stimulating electrode are visible in image on left and right sides, respectively. Aii. Individual inhibitory neurons express mDlx-jGCaMP8f and form axonal projections onto CA1 pyramidal cells within the stratum pyramidale and stratum radiatum, imaged at 40x magnification. B. Representative traces plotting the time-dependent change in jGCaM8f fluorescence relative to the total fluorescence (ΔF/F) imaged at 100 Hz at 40x magnification. Values were quantified from mDlx-jGCaMP8f-expressing projections within acute hippocampal slices. Ionotropic glutamate receptors were blocked via the addition of 5μM CNQX and 50μM AP5 in the bath. GABA_B_ receptors were blocked by 1 (red) or 10 μM (blue) CGP 55845. Bi. Scatter plot of subtracted peak amplitudes elicited by the second or fifth stimulus, compared to the first stimulus, in the train at 2Hz. Displayed P-values tabulated via paired two-tailed Student’s t-test. The experiment was conducted six times for six biological replicates. Bii. Scatterplot of the relative ratio of peak heights for each subsequent stimulus to the first at a frequency of 2 Hz. Connecting line plotted at arithmetic mean of each genotype. Displayed p-values refer to the effect of repetitive stimulation on jGCaMP8f transients and are tabulated as in Figure 1D. C. Representative electrically evoked jGCaMP8f traces elicited by high-frequency 10Hz stimulation in the presence or absence of 1 (red) or 10 μM (blue) CGP 55845. Ci. Scatterplot of subtracted peak amplitudes elicited by the second or fifth stimulus, compared to the first stimulus in the train, at a stimulus frequency of 10Hz. Cii. Scatterplot of the relative ratio of peak heights for each subsequent stimulus to the first at a frequency of 10 Hz. D. Representative trace of evoked mDlx-jGCaMP8f transients subsequent to the application of 2 depolarizing stimuli at 2 Hz. Traces containing 10μM baclofen are shown in green. Di. Scatter plot of the first or second amplitude in the presence or absence of 100nM, 1μM, or 1 μM baclofen. P-values shown tabulated by paired two-tailed Student’s t-test. Error bars represent means + S.E.M.

Stimulus-evoked responses of fluorescent amplitude plateaued by the third 10 ms frame post-stimulus (Figure 2B). At 2Hz, the amplitude of the second jGCaMP8f transient in the train was 120.7±6.5% of the first amplitude, while the fifth transient was 121.2±5.1% of the amplitude of the first (Figure 2Bi). In contrast, at 10 Hz (Figure 2C), the amplitude of the second and fifth transients in the train were reduced to 87.5±4.9% and 80.8±4.9%, respectively, of the first amplitude (Figure 2Ci). These experiments were then repeated in the presence of GABA_B_ receptor antagonist CGP 55845. In contrast to the effects of CGP 55845 on synaptic transmission evoked by high frequency stimulation as measured by IPSC amplitudes, there was no effect on the amplitudes of evoked presynaptic Ca^2+^ signals at 2 Hz (Figure 2Bii) or 10 Hz (Figure 2Cii). Treatment with CGP 55845 did not alter fluorescent transient peak heights for any stimulus (Supplemental Figure 3).

In contrast, application of 1µM baclofen inhibited evoked Ca^2+^ transients to a mean amplitude of 68.14± 18.3% of baseline, and 10 µM baclofen further to 43.95±7.7% of baseline (Figure 2D-2Di). This reduction in Ca^2+^ entry is consistent with the observed reduction of neurotransmitter release in baclofen in which IPSC amplitudes were reduced to a larger extent (Figure 1I), as there is a 3^rd^ to 4^th^ power relationship between Ca^2+^ entry and resultant synaptic transmission ^37^. This effect of baclofen is consistent with results at excitatory terminals of CA3 and CA1 pyramidal neurons, as well as observations at the calyceal synapse of Held and in cerebellar excitatory terminals. Results from both IPSC and fluorescence recordings are consistent with an inhibitory effect of baclofen-driven GABA_B_ receptor activation on GABA release caused by inhibition presynaptic Ca^2+^ influx. However, this effect on Ca^2+^ was not seen following repetitive evoked release, suggesting an alternate downstream target for endogenously released GABA acting at autoreceptors.

### Autoinhibitory GABA_B_ receptor activation by high frequency stimulus does not inhibit presynaptic Ca^2+^ signals

Epifluorescence imaging was centered on axo-somatic synapses in CA1. Nevertheless, the approach does not completely eliminate fluorescence of dendritic structures that may contaminate the recording. To overcome this, we used lattice light sheet microscopy to selectively illuminate single planes in volumes of hippocampal slices to isolate single jgCaMP8F labeled presynaptic terminals that surround the cell bodies of pyramidal neurons. Single lattice light planes, measuring 0.40 μm deep with an area of 52 × 52 µm area within the CA1 somatodendritic field (Fig. 3A), were imaged at a depth of up to 50 µm from the upper surface of the hippocampal slice. Fluorescence transients from axonal varicosities expressing jgCaMP8f were evoked by a stimulation protocol as shown in Figure 1A, in the presence of glutamate receptor inhibitors 5µM CNQX and 50µM AP5 in the bath solutions. The first evoked jgCaMP8f fluorescence transient in the train had an amplitude of 7.83±0.83% ΔF/F (Figure 3B). During 2 Hz stimulation, no reduction in the amplitude of subsequent fluorescent transients were observed (Figure 3C-E). Conversely, during high-frequency stimulation at 10 Hz (Figure 3F), the second and fifth transient amplitudes were once again lower, being reduced to 73.4±6.3% and 75.1±4.2% of the first, respectively (Figure 3G-3H). Thus, results from individual axonal varicosities were similar to those obtained in 40x epifluorescence imaging of stratum radiatum, in which no reduction in Ca^2+^ transient amplitude was evoked by repetitive stimulation at 2 Hz, while a small reduction was seen at 10 Hz. Taken together, we conclude that the mechanism by which endogenously released GABA mediates presynaptic inhibition via GABA_B_ autoreceptors involves a separate intracellular pathway from baclofen-driven VGCC inhibition.

**Figure 3:**
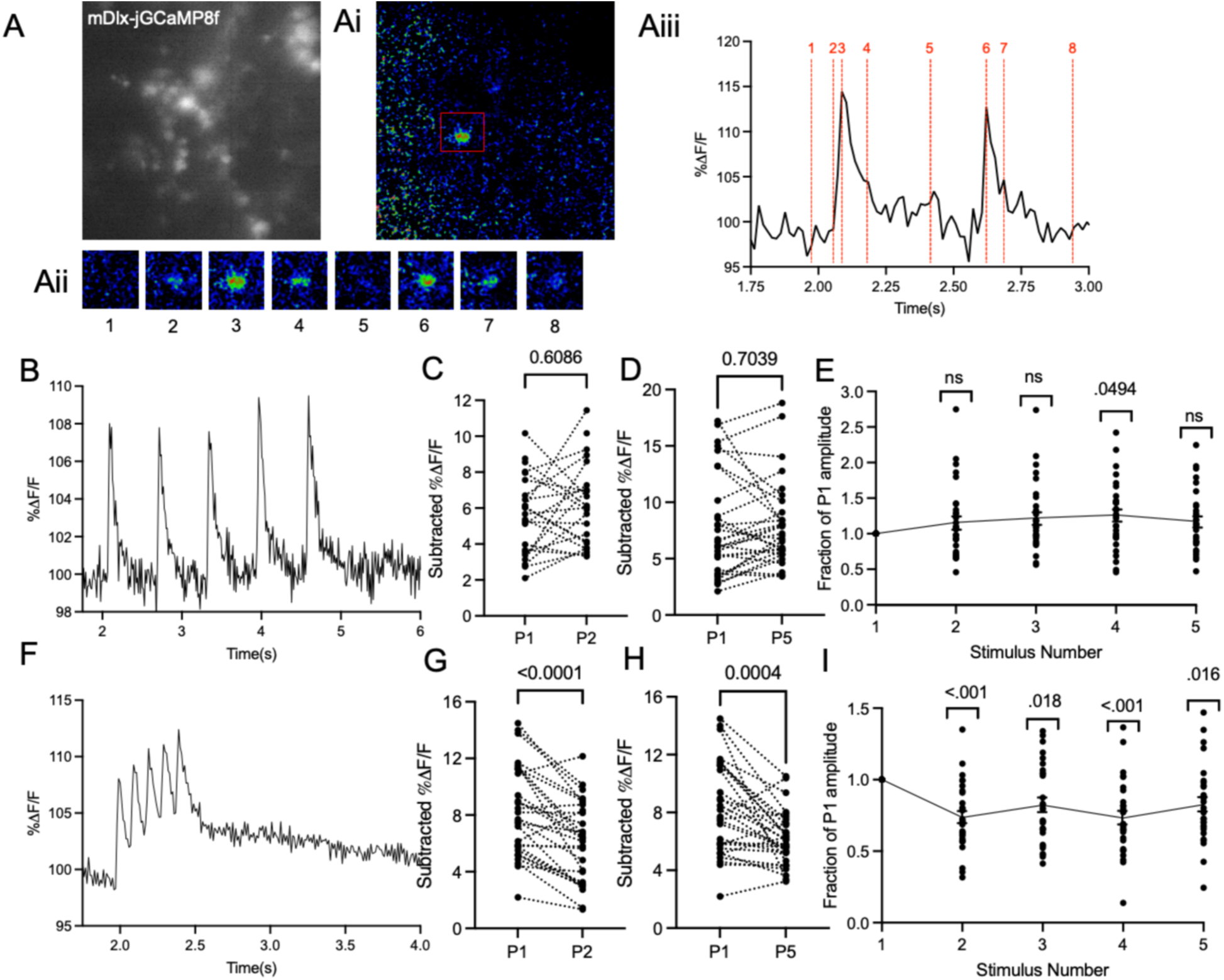
GABA_B_ autoreceptor activity localized to axonal projections from hippocampal inhibitory neurons. A. Image of single field of mDlx-jGCaMP8f-positive puncta at and adjacent to CA1 stratum pyramidale and radiatum, imaged under LLSM paradigm for recording and stimulation of acute *ex vivo* tissue. Ai. While imaging labeled GABAergic axons, application of depolarizing stimuli produced fluorescent transients along axonal varicosities (red box). False-color heatmap showing focal puncta of increased brightness subsequent to stimuli. Aii. Individual frames showing the time-dependent change in fluorescence of the region of interest in red in Ai. Aiii. Trace plotting the time-dependent change in jGCaM8f fluorescence relative to the total fluorescence (dF/F), imaged at 62 Hz. Numbers and dashed vertical lines in red denote individual frames within Aii. B. Representative trace from one singular puncta subsequent to five stimuli administered at 2 Hz. C. Scatterplot of the subtracted peak heights for the second stimuli in the train compared to the first. D. Scatterplot of subtracted peak heights for the fifth stimuli compared to the first. P-values on graph tabulated via paired two-tailed Student’s t-test. The results from 22 responding puncta from seven slices from five animals were included in the analysis. E. Scatterplot of the relative ratio of subtracted peak heights for each subsequent stimulus to the first at a frequency of 2 Hz: displayed p-values tabulated via two-way ANOVA with repeated measures with Bonferroni’s multiple comparisons test. A minor, yet significant rise was observed for the fourth stimuli in the train. F. Representative trace from one singular puncta showing suppression later in the train subsequent to five high-frequency stimuli administered at 10 Hz. G. Scattterplot of subtracted peak heights for the second stimuli in the train compared to the first. H. Scatterplot showing the subtracted peak heights for the fifth stimuli compared to the first. I. Scatterplot of the relative ratio of subtracted peak heights for each subsequent stimulus to the first after high-frequency stimulation. Error bars represent means + S.E.M.

### Autoinhibitory loss of GABA signaling is mediated by direct Gβγ-SNARE complex interaction

Despite a lack of observed effect on presynaptic Ca^2+^ entry by 2 or 5 Hz trains, the GABAergic autoreceptor pathway is clearly capable of inhibiting neurotransmission. Therefore, we sought to identify other mechanisms through which the presynaptic autoreceptors could reduce IPSC amplitudes to CA1 pyramidal cells. One possible mechanism is the Gβγ-SNARE pathway. Upon activation of a Gα_i/o_-type GPCR, liberated Gβγ subunits bind to the C-terminus of SNAP25^38,39^ within SNARE complexes and competitively inhibit synaptotagmin I binding and subsequent neurotransmitter release^40,41^. To determine whether this mechanism contributes to autoreceptor signaling in hippocampus, we utilized a homozygous SNAP25Δ3 mouse model, in which the *Snap25* locus carries the G204* mutation. This mutation causes reduced binding of SNAP25 to Gβγ^1,42^, but no change in overall basal exocytosis.

Acute hippocampal slices were prepared from adult male SNAP25Δ3 homozygotes or littermates carrying the wild-type *Snap25* allele, and CA1 pyramidal cells were whole cell patch clamped as in Figure 1. Electrically-evoked IPSCs were recorded in CNQX and AP5 as in Figure 1 and trains of stimuli were given at 2, 5, 10, and 20 Hz (Figure 4). We observed no effect of *Snap25* genotype on the recorded amplitudes of the first response in the train using a fixed stimulation amplitude of 50 µA and 200 μs (Figure 4A). In SNAP25WT mice, we observed significant IPSC suppression later in subsequent IPSCs during the 2 Hz train, consistent with non-transgenic animals in Figure 1. The second response in the train was 62.4±3.3% of the first, dropping to 56.7±3.3% for the fifth event (Figure 4Ai-4Aii). In contrast, the SNAP25Δ3 genotype significantly attenuated the autoreceptor suppression, with the second IPSC amplitudes reduced only to 86.7±6.4%, and the fifth reduced to 81.5±5.97 (Figure 4Aiii). At a stimulus frequency of 5 Hz, the effect of genotype was still present, but only significant for the final stimuli in the trains (Figure 4B-4Bi). The effect of genotype was more prominent during high-frequency 10 Hz stimulation, where the second WT IPSC amplitude was 63.4±4.4% of the first, and the fifth 52.3±4.625 (Figure 4C-4Ci). In comparison, the second SNAP25Δ3 IPSC amplitude was 87.5±8.1% of the first, and the fifth 76.4±5.68%. A similar trend was observed at a stimulus frequency of 20 Hz (Figure 4D-4Di). The rise time or decay time of the IPSC amplitudes within the paradigm did not differ by genotype (Supplemental Figure 4). From this, we conclude that GABA_B_ autoreceptors suppression of GABA release is strongly attenuated during stimulus trains in SNAP25Δ3 homozygotes. These results suggest that GABA_B_ autoreceptor suppression acts via the Gβγ-SNARE interaction, in all cases and concomitantly with its inhibition of presynaptic Ca^2+^ channel activity, particularly late in stimulus trains.

**Figure 4:**
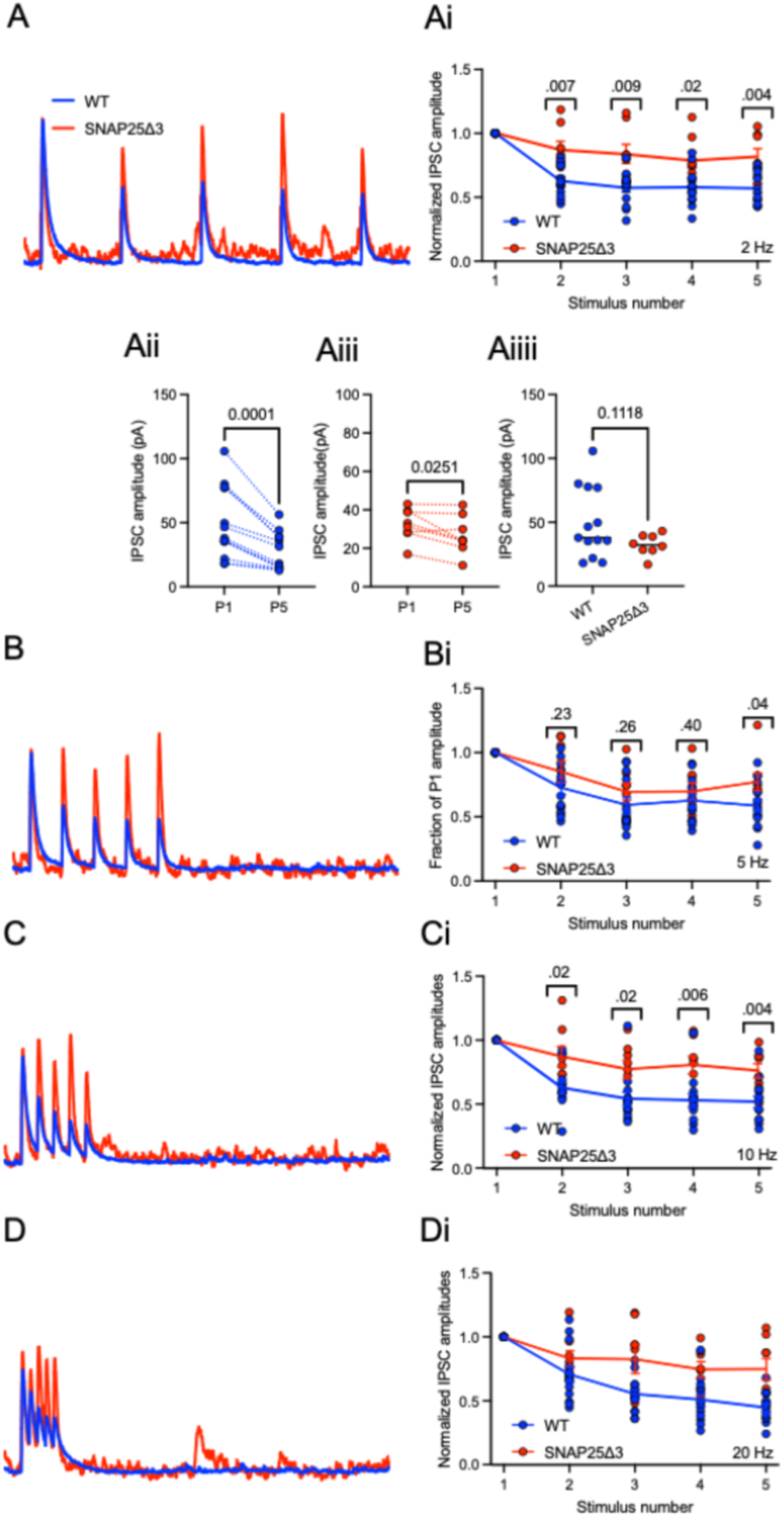
GABA_B_ autoreceptors signal via the Gβγ-SNARE mechanism. A. Representative traces of IPSCs subsequent to a train of five stimuli in hippocampal slices from WT animals (blue) or SNAP25Δ3 littermates (red) at 2 Hz. Ai. Scatterplot of the relative ratio of peak amplitudes for each subsequent stimulus to the first in the train at a frequency of 2 Hz. Connecting line plotted at arithmetic mean of each genotype. The experiment was performed up to 13 times with at least ten WT biological replicates, and eight SNAP25Δ3 biological replicates. Error bars represent means + S.E.M. Aii. Scatterplot of peak IPSC amplitudes for the fifth stimulus in the train compared to the first stimulus in the train in WT CA1 pyramidal cells. Aiii. Scatterplot of peak IPSC amplitudes for the fifth stimulus in the train compared to the first stimulus in the train in SNAP25Δ3 CA1 pyramidal cells. Aiii. P-values on graphs tabulated via paired two-tailed Student’s t-test. Aiiii. Scatterplot comparing peak IPSC amplitudes for the first stimulus in the train for WT and SNAP25Δ3 cells. P-values on graph obtained via unpaired two-tailed Student’s t-test. B. Representative traces of IPSCs as in (A) at a stimulus frequency of 5 Hz. Bi. Scatterplot of the relative ratio of peak amplitudes as in (Ai) at 5 Hz. C. Representative traces of IPSCs as in (A) at 10 Hz. Ci. Scatterplot of the relative ratio of peak amplitudes as in (Ai) at 10 Hz. D. Representative traces of IPSCs as in (A) at 20 Hz. Di. Scatterplot of the relative ratio of peak amplitudes as in (Ai) at 20 Hz.

### GABAergic synapses in SNAP25Δ3 hippocampus exhibit altered suppression of GABA release

Autoreceptors act to reduce neurotransmitter release from presynaptic terminals during stimulus trains, but changes to IPSC amplitudes might also result from signaling events within the postsynaptic CA1 pyramidal cell. To isolate the effect of autoreceptors on GABA release from inhibitory neurons in CA1 dendritic fields, WT and SNAP25Δ3 animals were transduced with the genetically encoded GABA biosensor, iGABASnFr2^30,43^ under the control of the pan-neuronal promoter *syn*, via stereotaxic AAV injection into the hippocampus (Figure 5A). iGABASnFr2 fluorescence transients were then electrically evoked in acute hippocampal slices using the same stimulation protocols as those for electrophysiological recordings and recorded via epifluorescence microscopy as in Figure 2. Basal iGABASnFr2 fluorescence was uniform throughout the CA1 stratrum radiatum (Figure 5Ai-Aii). Trains of monosynaptic iGABASnFr2 responses were obtained at 2 Hz (Figure 5B), with slower off rates compared to jgCaMP8f. At 2 Hz, a decay of the iGABASnFr2 signal (shaded in purple) was observed in slices from WT animals, but in slices from SNAP25Δ3 animals, iGABASnFr2 fluorescence showed continuous accumulation throughout the pulse train. At 2 Hz, the second response was inhibited to 63.1±5.16% of the first in WT and 64.5±3.76% in SNAP25D3 (Figure 5Bi). However, the total magnitude of iGABASnFr2 fluorescence was significantly larger in SNAP25Δ3 (Figure 5Bii) after 2 and 5 stimuli. At 10 Hz, the second response was inhibited more completely, to 30.1±1.93% of the first in WT, but significantly less so in SNAP25Δ3, to 57.8±7.7% (Figure 5Ci), and the total magnitude of iGABASnFr2 fluorescence significantly accumulated much more in SNAP25Δ3 (Figure 5Cii). Suppression of iGABASnFr2 transients on the second and fifth stimulus were observed in both WT and SNAP25Δ3 slices (Figure 5D). No effect of genotype was observed for individual release events within the train (Supplemental Figure 5). Time constants of decay were also significantly larger in SNAP25Δ3 slices than WT slices (Supplemental Figure 6). While the sensor fluorescence kinetics are too slow to fully recapitulate the rapid kinetics of GABA release, it does resolve a substantial difference between stimulus train-evoked GABA release in WT vs SNAP25Δ3. Thus, our stimulus paradigm is sufficient to activate GABA_B_ autoreceptors on the presynaptic terminals of hippocampal inhibitory neurons, and reveals that the functional effect of these receptors is to suppress further release of GABA into the cleft: this effect is diminished in SNAP25Δ3 during high-frequency stimulation.

**Figure 5:**
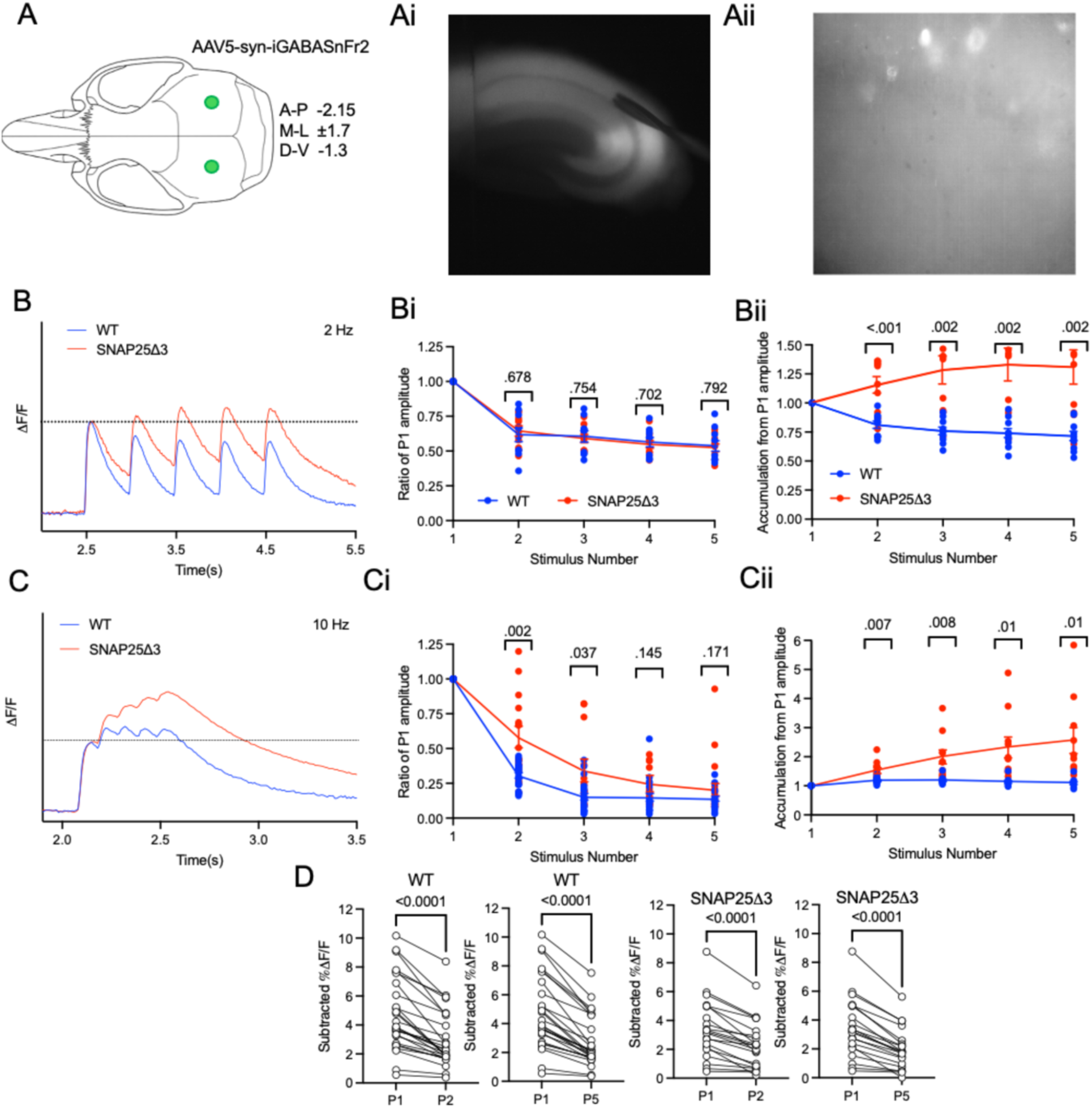
Presynaptic GABA release is regulated by GABA_B_ autoreceptors and the Gbg-SNARE pathway. A. Visual schematic of stereotactic injection of AAV1-syn-iGABASnFr2-WPRE into mouse parenchyma. Ai. Acute mouse hippocampal slice expressing uniform syn-iGABASnFr2 fluorescence imaged via epifluorescence microscopy at 4x magnification. Retaining harp and stimulating electrode are visible in image on left and right sides, respectively. Aii. Fields of neuropil within CA1 stratum radiatum express uniform syn-iGABASnFr2 fluorescence of moderate brightness at 40x magnification. B. Averaged trace showing the time-dependent change in iGABASnFr2 fluorescence relative to the total fluorescence (ΔF/F) subsequent to the application of 5 depolarizing stimuli at 2Hz at t = 2.5 to 4.5 s, imaged at a rate of 100 Hz. Data recorded from slices from WT animals is plotted in blue, while data recorded from slices from SNAP25Δ3 animals is plotted in red. Areas above the initial peak are shaded in green, while areas below it are shaded in purple. Bi. Scatterplot of the relative ratio of subtracted peak heights for each subsequent stimulus to the first at a frequency of 2 Hz: displayed p-values tabulated via two-way ANOVA with repeated measures with Bonferroni’s multiple comparisons test. Bii. Scatterplot of the ratio of the accumulated total iGABASnFr2 signal for each subsequent stimulus relative to the first at a frequency of 2 Hz. Experiments were repeated ten times from eight independent biological replicates per genotype. C. Averaged trace as in Figure 5B showing the time-dependent change in iGABASnFr2 fluorescence after high-frequency stimulation at 10Hz. Ci. Scatterplot of the relative ratio of subtracted peak heights for each subsequent stimulus after the first at 10 Hz as in Fig. 5C. Cii.. . Scatterplot of the ratio of the accumulated total iGABASnFr2 signal for each subsequent stimulus after the first at a frequency of 10 Hz. Experiments were repeated 21 times from at least eight biological replicates per genotype. D. Scatterplot showing subtracted dF/F values for the first, second, and fifth peaks in the train for WT or SNAP25Δ3 slices. Displayed P-values tabulated via paired two-tailed Student’s t-test. Error bars represent means + S.E.M.

### Prevention of presynaptic Gβγ-SNARE interaction by SNAP25Δ3 blocks inhibitory effects of GABA_B_ receptor activity on mIPSCs

Our data indicate that activation of GABA_B_ receptors on axons and presynaptic terminals of GABAergic neurons leads to Gβγ interaction with both Ca^2+^ channels and directly with the SNARE complex, whereby the downstream target depends on the method of GABA_B_ activation. In repetitive evoked responses, released GABA acts at autoreceptors to inhibit release at the SNARE complex. In contrast, baclofen inhibits Ca^2+^ entry through VGCCs. This may result from either highly localized active zone receptor activation, or more dispersed receptor activation via reduced cytosolic cAMP concentrations. Resulting changes in Ca^2+^ entry are expected to change downstream mIPSC frequency. However, in contrast, changes in synaptic strength following Gβγ interaction with the SNARE complex have not been shown to alter release probability at excitatory synapses but did reduce single evoked quantal amplitudes^44^ and the concentration of neurotransmitter released into individual synaptic clefts^39,45,46^. We hypothesized that if GABA_B_ receptors reduced presynaptic GABA release from inhibitory neurons, both mIPSC frequencies and amplitudes would fall correspondingly in the presence of GABA_B_ agonists, and that this effect would be reduced in SNAP25Δ3 homozygous mice. To test this, we applied baclofen while recording mIPSCs from CA1 pyramidal neurons from WT and SNAP25Δ3 slices (Fig 6A-6B). Neurons were recorded under whole cell conditions, with pipettes containing 140 mM of Cl^-^ to ensure that GABA_A_ receptor mediated currents were inward at resting membrane potentials. The tissue was superfused with the Na^+^ channel blocker tetrodotoxin (TTX 1µM) to prevent synaptic activity caused by action potentials, and with CNQX and AP5 to block spontaneous glutamatergic events.

**Figure 6.**
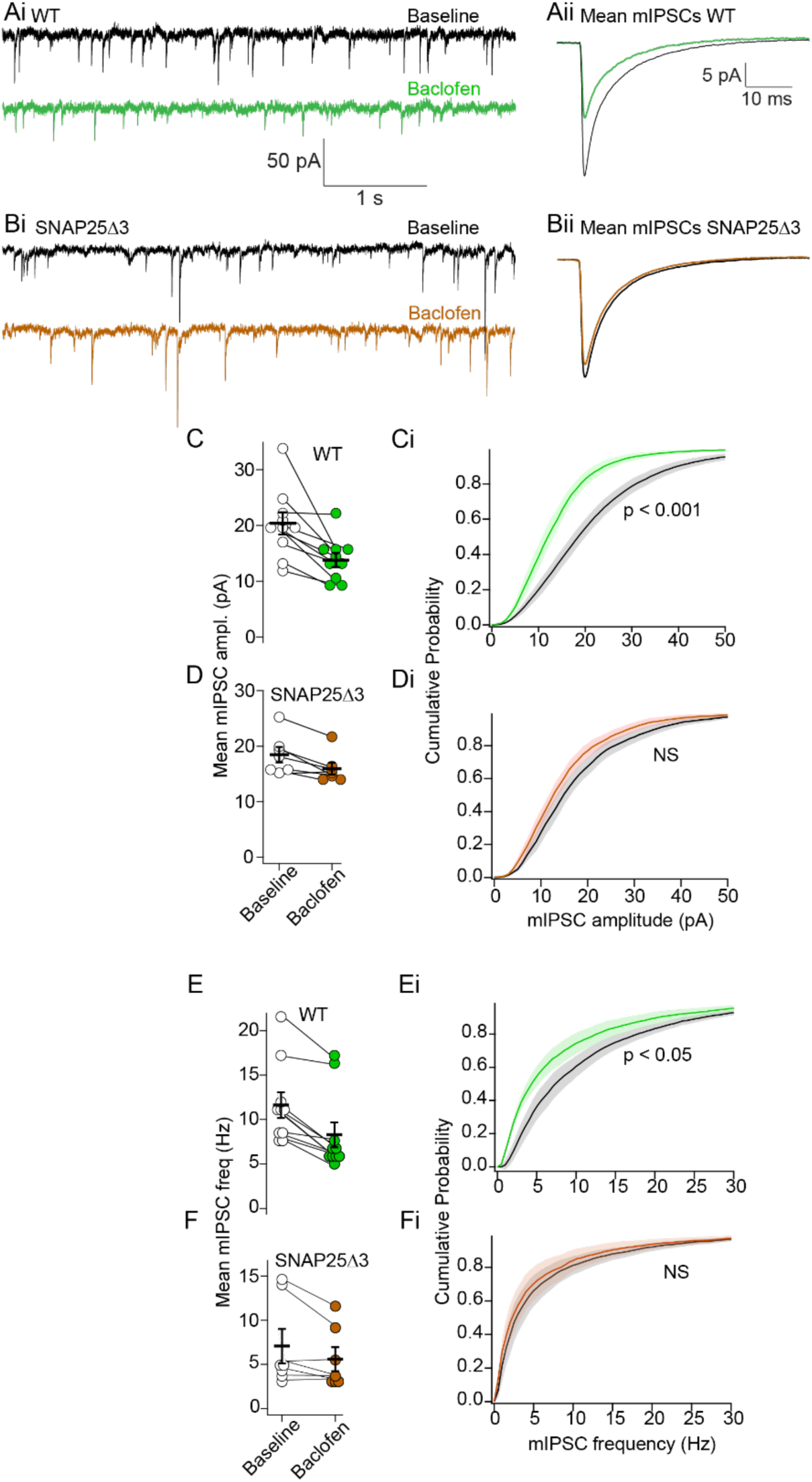
GABA_B_ signals via the Gβγ-SNARE mechanism to reduce mIPSC amplitudes and frequencies. Ai. Representative trace of miniature inhibitory postsynaptic currents (mIPSCs) recorded from WT CA1 pyramidal cells within acute slices in the presence of 1μM tetrodotoxin(TTX), 5μM CNQX and 50μM AP5. Baseline recordings are shown in black, while recordings in the presence of 10mM baclofen are shown in green. Aii. Averaged trace of individual mIPSC event from WT cells in the presence or absence of baclofen. Bi. Representative trace of mIPSCs recorded from SNAP25Δ3 cells. Baseline recordings are shown in black, while recordings in the presence of 10mM baclofen are shown in brown. Bii. Averaged trace of individual mIPSC event from SNAP25Δ3 in the presence or absence of baclofen. C. Scatterplot of mean mIPSC amplitudes in the presence or absence of baclofen. Baclofen significantly reduced mIPSC amplitudes in WT. Displayed P-values tabulated via paired two-tailed Student’s t-test. Error bars represent means + S.E.M. Ci. Cumulative distribution plot of mIPSC amplitudes from WT neurons in the presence or absence of baclofen (green). Displayed P-values for effect of baclofen on the distribution of amplitudes was tabulated by the two-sample Kolmogorov-Smirnov test. . D. Scatterplot of mean mIPSC amplitudes in SNAP25Δ3 neurons in the presence or absence of baclofen (brown). Di. Cumulative distribution plot of mIPSC amplitudes from SNAP25Δ3 neurons in the presence or absence of baclofen. E. Scatterplot of mean mIPSC frequencies in the presence or absence of baclofen(green). Baclofen correspondingly significantly reduced mIPSC frequencies in WT. Ei. Cumulative distribution plot of mIPSC frequencies from WT in the presence or absence of baclofen. F. Scatterplot of mean mIPSC frequencies in SNAP25Δ3 neurons in the presence or absence of baclofen(brown.) Fi. Cumulative distribution plot of mIPSC frequencies from SNAP25Δ3 in the presence or absence of baclofen.

We measured amplitudes and frequencies of the detected events and simultaneously monitored whole cell access and membrane integrity (series resistance, Rs and membrane resistance Rm) with a 5mV negative potential pulse at 30 s intervals. Neither Rs nor Rm varied significantly. Mean Rs at baseline was 20.1 ± 1.3 MΩ and 20.4 ± 1.8 MΩ in baclofen (n = 9, p = . Mean Rm was 106 ± 23 MΩ at baseline and 97 ± 11 MΩ in baclofen. Similarly, holding currents were unaffected by baclofen application (holding current at a membrane potential of -60 mV, -79 ±37 pA mV at baseline, -99 ± 36 pA in 10 µM baclofen, n=10, p = 0.15, t(9) =2.2) .

Both mIPSC amplitudes and event frequencies were substantially reduced by baclofen in neurons recorded from WT animals (Figure 6A). Mean event amplitudes were reduced to 65 ± 5% of baseline: from 21.4 ±1.7 pA at baseline to 13.9 ± 1.3 pA in baclofen (Figure 6Ai–Figure 6C). Mean frequencies were reduced to 70.5 ± 0.5 % of the baseline: from 11.4 ± 1.4 Hz at baseline to 8.2 ± 1.4 Hz in baclofen (Figure 6E). The profound effect on mIPSC amplitude by baclofen is typically assumed to result from a postsynaptic effect^47^; however, there was no effect of baclofen on Rm or on holding current. Indeed, the neurons were recorded for at least 20 mins in whole cell conditions prior to experimental recordings to allow TTX and glutamate antagonist equilibrium, which will eliminate many intracellular signaling pathways in the postsynaptic neuron. However, Gβγ targeting of the SNARE complex is known to directly reduce synaptic cleft concentration of neurotransmitter during evoked synaptic transmission in glutamatergic synapses in lamprey^44,46,48^, chromaffin cells in culture^49,50^ , and in hippocampal pyramidal neurons^24,45^.

Thus, we used the same experimental approach in slices obtained from SNAP25Δ3 mice (Figure 6B). In these recordings, event amplitudes mediated by baclofen were reduced from 18.4 ± 1.9 pA to 15.5 ± 1.4 pA (to only 85 ± 3%) of baseline amplitudes, and mean frequencies were reduced by 70.5 ± 0.5 %, shifting from 11.4 ± 1.4 Hz at baseline to 8.2 ± 1.4 Hz in baclofen (Figure 6Bi, 6D, 6F, Supplemental Figure 7). Taken together, we infer that the GABA_B_ receptor reduces mIPSC event sizes and frequencies through the Gβγ-SNARE pathway, reinforcing our hypothesis of a presynaptic mechanism for GABA_B_ in hippocampal inhibitory neurons.

### cAMP signaling pathways govern GABA_B_ receptor function

We have demonstrated that presynaptic GABA_B_ receptors inhibit neurotransmission by acting as autoreceptors which release Gβγ to target the SNARE complex directly. This target is activated by rapid endogenous release of GABA, while presynaptic evoked Ca^2+^ entry is unaffected. However, when the exogenous GABA_B_ receptor agonist, baclofen, is added to the superfusate it potently inhibits presynaptic Ca^2+^ entry.

While direct binding of Gβγ subunits to Ca_V_2 channels (N, P/Q, and R-type) channels is understood as the canonical mechanism for reduction of Ca^2+^ entry, these Ca_V_2 channels are also regulated through cAMP-dependent pathways such as PKA^51–54^ phosphorylation. Phosphorylation of the channel may raise or lower channel conductances or kinetics. We hypothesized that with the longer time frames of minutes that it takes to add baclofen to the superfusate, baclofen would reduce cytosolic cAMP levels by liberation of Gα_i_ or perhaps Gβγ subunits to bind to and inhibit adenylyl cyclase, which would then alter the activity of one or more cAMP effectors such as PKA or HCN channels. If this is the case, raising the activity of adenylyl cyclase with forskolin^55^ should greatly diminish the effects of baclofen. To test this, IPSCs were recorded from CA1 pyramidal cells using patch clamp, in the presence of 50 µM forskolin for a minimum of 10 minutes. Application of forskolin alone did not change individual isolated IPSC amplitudes (Figure 7A-7Ai). Next, the intermediate and saturating concentrations of baclofen, 1 µM and 10 µM, were applied and IPSCs were recorded once more. Without forskolin, 1 µM baclofen significantly reduced IPSC amplitudes to 25.6 ± 3.69 % of control amplitudes while 10 µM baclofen reduced amplitudes further to 15.0 ± 3.53 % of control amplitudes (Figure 7Aii). In contrast, in slices pretreated with forskolin, 1 µM baclofen did not significantly reduce IPSC amplitudes (74.4 ± 14.0 % of control amplitudes), while 10 µM baclofen significantly reduced IPSC amplitudes to 16.8 ± 6.73 % of control amplitudes (Figure 7Aiii). Given this effect of forskolin on IPSC amplitudes, we then sought to determine if forskolin pretreatment could similarly diminish the inhibitory effect of baclofen upon whole-neuron mDlx-jgCaMP8f transients using the same paradigm as Figure 4. Application of forskolin alone did not change the electrically evoked rise in the jGCaMP8f signal (Figure 7B) at 2 Hz, with peak amplitudes being 104.9±14.8% of control amplitudes. By comparison, the inhibitory action of baclofen on mDlx-jGCaMP8f Ca^2+^ transients was blunted significantly by forskolin. 1 µM baclofen reduced P1 amplitudes to only 91.2 ±13.2% of control amplitudes in the presence of forskolin, while 10 µM baclofen reduced P1 amplitudes to 74.3 ±14.6 % of control amplitudes (Figure 7Bi-7Bii). From this, we conclude that the action of baclofen on GABA_B_ in inhibitory neurons occurs partially through a reduction in inhibitory neuronal cytosolic cAMP levels.

**Figure 7:**
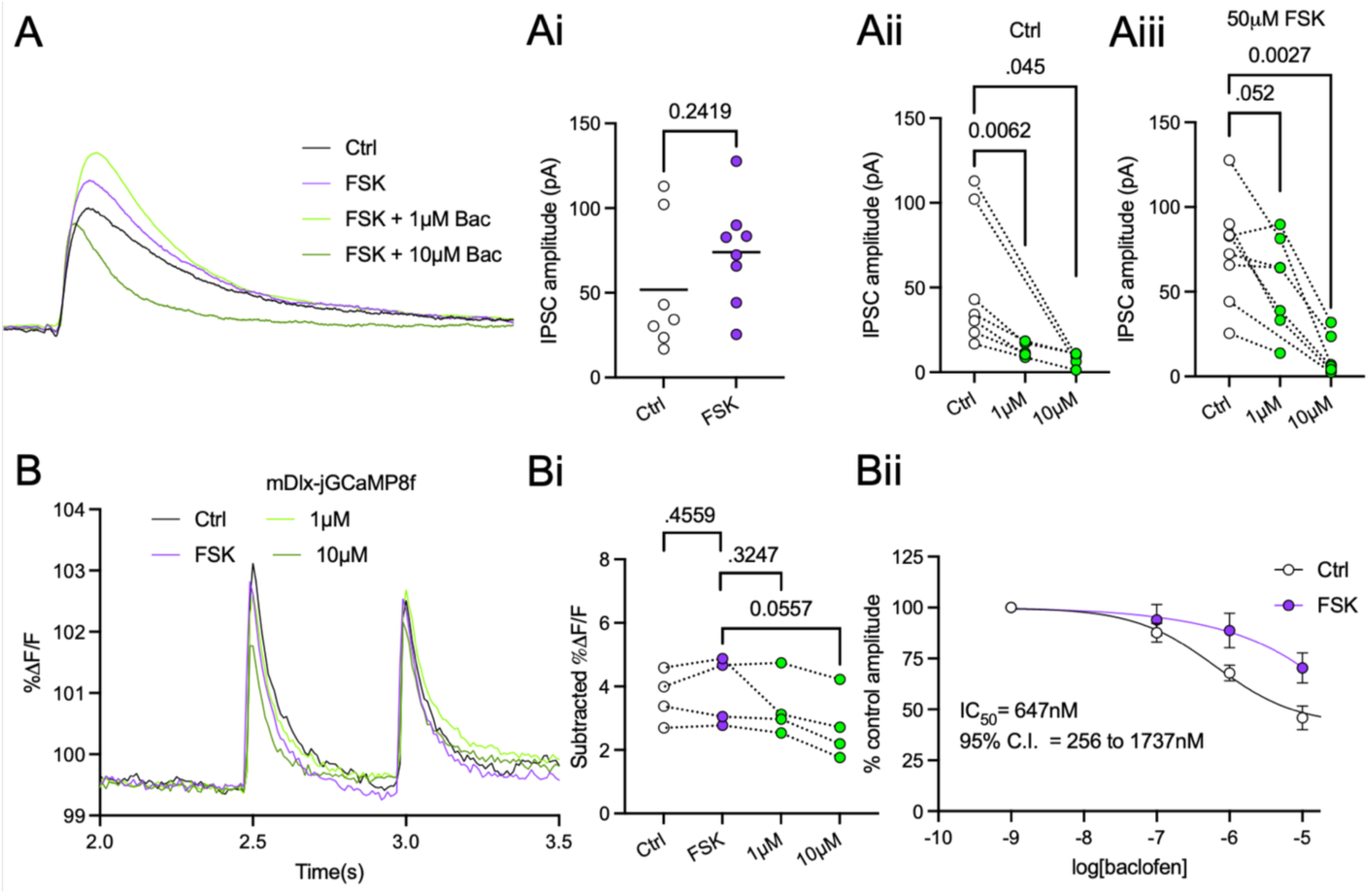
Baclofen signals through GABA_B_ in part via inhibition of adenylyl cyclase. A. Representative trace of IPSCs in the presence or absence of forskolin (purple) or forskolin + baclofen (green). Ai. Scatterplot of electrically evoked IPSCs from CA1 pyramidal neurons held the whole-cell voltage-clamp configuration as in Figure 1, in the tabulated in the presence (white) or absence (purple) of 50 μM forskolin administered in the bath. Displayed P-values tabulated via unpaired two-tailed Student’s t-test. Aii. IPSC amplitudes from slices not incubated with forskolin, treated with 1 or 10μM baclofen in the bath. Data redisplayed from Figure 1F. Displayed P-values tabulated via paired two-tailed Student’s t-test. Aiii. IPSC amplitudes as in Figure 6Ai, from slices treated with forskolin. B. Representative trace from slices virally transduced with mDlx-jGCaMP8f as in Figure 2 showing the time-dependent change in electrically evoked fluorescence relative to the total fluorescence (dF/F) subsequent to the application of 2 depolarizing stimuli at 2 Hz at t = 2.5 and 3 s, imaged at a rate of 100 Hz in the presence of CNQX and AP5. Traces containing forskolin but not baclofen are shown in purple, while traces containing both are shown in green. Bi. Scatterplots of subtracted jGCaMP8f peak heights for each condition depicted in Figure 6B. Displayed P-values tabulated via paired two-tailed Student’s t-test. Bii. Sigmoidal concentration-response relationships for the effect of baclofen on jGCaMP8f peak heights in the presence and absence of forskolin. Error bars represent means + S.E.M.

## DISCUSSION

GABA_B_ autoreceptors have long been considered ubiquitous at cortical GABAergic presynaptic terminals. We illustrate a new role for the Gβγ-SNARE pathway in this autoreceptor function within hippocampal inhibitory neurons. Three principal targets of membrane delimited Gβγ effects that rapidly control neurotransmitter release have been proposed at presynaptic terminals. These are: (1) Inhibition of Ca^2+^ channels by a direct action of Gβγ leading to reduced presynaptic Ca^2+^ entry and a lowered probability of neurotransmitter release. (2) The activation of G protein inwardly rectifying K+ (GIRK) currents. GIRK current activation will alter presynaptic Ca^2+^ entry. The application of an exogenous agonist, baclofen, has shown that GABA receptors can inhibit presynaptic Ca^2+^ entry^24,25,27,56–58^, and this finding is reinforced here through the effect of baclofen on electrically evoked concentration-dependent inhibition of mDlx-jgCaMP8f transients **(Fig. 2)**. (3), Gβγ signaling downstream of Ca^2+^ entry results from Gβγ acting directly at the SNARE complex. This action of Gβγ is downstream from Ca^2+^ entry and we now demonstrate that GABA autoreceptors that effect the same GABA release, but that are activated by physiologically relevant GABA release rather than exogenous baclofen do not alter presynaptic Ca^2+^ entry, but rather they act by a direct Gβγ-SNARE interaction to control further GABA release.

The classical view of GABA_B_ autoreceptors has been that GABA release from presynaptic terminals inhibits further release by a rapid effect of released membrane bound Gβγ acting to inhibit presynaptic Ca^2+^ entry on following repeated action potential firing. it has been thought that these autoreceptors act solely by inhibiting presynaptic Ca^2+^ entry through VGCCs. Indeed, in the hippocampus, presynaptic Ca^2+^ entry is potently inhibited by the GABA_B_ receptor agonist, baclofen, at excitatory termimals in both CA1^25^ and subiculum^24,45^ and we now show a similar result for inhibitory presynaptic terminals.

While GABA_B_-mediated Gβγ release activates hyperpolarizing currents via GIRK channels at the postsynaptic density^20,22,23,47^, a role for GIRK at presynaptic terminals has been less well-supported. It was reported that no presynaptic GIRK currents are activated by GABA_B_ receptors^21,26,47,59^ in multiple neuronal populations, while other groups report that the GIRK channel blocker tertiapin-Q was able to reverse the inhibitory action of GABA_B_ on chemically evoked glutamate release in isolated cortical synaptosomes^56^. Axonal GIRK conductances have been reported to occur at hippocampal GABAergic neurons^47^. If presynaptic GIRK conductances are activated by baclofen, then inhibition would act by subsequent reduced Ca^2+^ entry.

The action of Gβγ directly at SNARE complexes was first identified in glutamatergic reticulospinal neurons in lamprey^41^ and has since been identified in PC12 cells in culture^24,40,60^, hippocampal CA1 and CA3 pyramidal neurons, and in amygdala, nucleus accumbens^28,61,62^ and Chromaffin cells^50^ . The development of a mutant mouse lacking this site^1^ enabled screening for this effect in a number of studies. A role for Gβγ-SNARE in autoreception has been demonstrated in postganglionic sympathetic innervation of adipose tissue^63^ and cone ribbon synapses of the retina^64^ in salamander. It remains unclear whether the mGluR7 glutamatergic autoreceptors that regulate excitatory neurotransmission from CA3 pyramidal cells onto CA1 signal via the Gβγ-SNARE mechanism as inhibitory GABA_B_ autoreceptors function.

GABA_B_ receptors, whether acting as heteroreceptors or autoreceptors, have been shown to act by multiple mechanisms including inhibition of Ca^2+^ entry^25^ as well as mechanisms downstream from Ca^2+^ entry^65^. In glutamatergic afferents to both PV+ inhibitory neurons and medium spiny neurons in the nucleus accumbens, GABA_B_ heteroreceptors drive presynaptic inhibition via the Gβγ-SNARE pathway^28,62^. We now show that in evoked responses GABA_B_ autoreceptor effects are markedly reduced in SNAP25Δ3 mice while block of GABA_B_ receptors has no effect on repetitive stimulation evoked presynaptic Ca^2+^ transients. Nevertheless, during high-frequency stimulation (Figure 2-3), GABA concentrations are higher, and more likely to spill out of the synapse, activating extrasynaptic receptors. We show that fast dynamic GABA_B_ signaling (ms to s) via the endogenous ligand GABA utilizes the Gβγ-SNARE pathway, but tonic equilibrium activation of GABA_B_ by exogenous agonists such as baclofen also works independently of the Gβγ-SNARE mechanism, since no effects of the SNAP25Δ3 mutation were observed (Figure 1, 3, 5). It seems likely that the inhibitory effects of baclofen on evoked GABA release preclude recording downstream effects on the SNARE complex. Indeed, without evoked release we observed considerable SNAP25Δ3 effects on mini frequency and amplitude.

In contrast, on the time scale of minutes to hours, one or more adenylyl cyclase-dependent pathways utilized by GABA_B_ contributes considerably to the large and uniform suppression of evoked Ca^2+^ entry observed at saturating concentrations of baclofen. IPSC suppression by baclofen could be bypassed via the addition of the adenylyl cyclase allosteric activator forskolin^66^, suggesting that baclofen may suppress GABA release via the inhibitory action of Gα_i/o_ on adenylyl cyclase (Figure 5). However, adenylyl cyclase is unlikely to cause dramatic changes in cAMP concentrations at the presynaptic terminal within milliseconds in response to released GABA. At equilibrium, baclofen was still able to inhibit mDlx-jgCaMP8f transients in the presence of forskolin, albeit to a significantly reduced extent. The inhibitory action of Gβγ on VGCC may be altered by channel phosphorylation^51,52^, though the number and nature of the exact cAMP-dependent effectors in the process remains largely unknown.

In this study, we used the recently developed fluorescent biosensor iGABASnFr2^30^ to confirm our results from conventional electrophysiological studies, especially that the effects of either baclofen or GABA_B_ autoreceptors activated by release of GABA, are indeed presynaptic. iGABASnFr2 has slower kinetics and is less sensitive than specific expression of jGCaMP8f in inhibitory neurons, but is needed to provide inferences for inhibitory mechanisms occurring downstream of Ca^2+^ entry, and can also measure post-release GABA dynamics directly. Reduction of mIPSC frequencies (Figure 6E-6F) would strongly suggest a presynaptic mechanism for GABA_B_, consistent with prior results. While inhibitory postsynaptic current amplitudes could change if pyramidal GABA_A_ channel insertion/internalization or other determinants of chloride conductance are altered by the recording paradigm, the use of iGABASnFr2 is not subject to these experimental caveats. However, the time resolution of electrophysiological studies is greatly superior to that of iGABASnFr2 due to the slow unbinding rates of GABA as well as binding rates still slower than the rate of diffusion. With a decay constant of ∼0.2 s, some GABA released during P1 is likely still bound to iGABASnFr2 at the time of the P2 stimulus at 2Hz: this forms the basis for the accumulation observed in Figure 5. While the measured decay constants are larger in SNAP25Δ3, the electrophysiological studies in Fig. 2 do not support a model in which alterations in GABA reuptake or degradation are the primary factors for the SNAP25Δ3 phenotype, since IPSC amplitudes return to unity between stimuli. Instead, any alterations in GABA reuptake kinetics in SNAP25Δ3 occur secondary to a presynaptic autoreceptor-based mechanism (Figure 5C).

The studies in this manuscript highlight the importance of kinetics of agonist application as a determinant of receptor function. The release of synaptic GABA on autoreceptor signaling is not phenocopied by the application of baclofen. The receptors activated by baclofen are likely to include both autoreceptors adjacent to the presynaptic terminal and discrete populations of extrasynaptic GABA_Β_ receptors endogenously activated by tonic GABA currents: the localization of these receptors may determine the signaling mechanism activated downstream of GABA binding and while baclofen may inhibit both at Ca^2+^ channels and the SNARE complex, its efficacy at inhibiting upstream Ca^2+^ likely dominates its measured effects during evoked release. In contrast during recording of mIPSCs baclofen’s actions will be on effects downstream of Ca^2+^ entry which likely explains the loss of effect in SNAP25Δ3 mice.

These GABA_B_ autoreceptor effects at the SNARE complex may be widespread. In cortex, an inhibitory action of GABA_B_ has been observed on mEPSC frequencies^65,67^ and mIPSCs^67,68^. Disruption of the Gβγ-SNARE pathway blocked the effect of baclofen on former, but not the latter^67^. Given that substantial similarities in gene expression exist between cortical and hippocampal inhibitory neurons of the class 06/07 CTX-MGE and CTX-CGE^69^, we anticipate that cortical inhibitory neurons should also utilize the Gβγ-SNARE mechanism for GABA_B_-dependent autoreceptor function, similar to the hippocampal inhibitory neurons shown here. Nevertheless, variations between regions stemming from differential local circuit activity, with resultant changes in gene expression driving recruitment of other G_i/o_-delineated signaling pathways.

In conclusion, we assessed the relative contributions of presynaptic Ca^2+^, presynaptic cAMP, and the contribution of Gβγ signaling at the SNARE complex downstream of both in GABA_B_ autoreceptor function. The mechanisms through which GABA_B_ receptors reduce GABA release are key determinants of excitatory/inhibitory balance, which is central to cellular and synaptic homeostasis. Any homeostatic change that increases excitatory neurotransmission or decreases inhibitory neurotransmission increases the probability of seizure occurrence^70^. A less completely understood example of E-I imbalance is thought to exacerbate neurodegeneration in Alzheimer’s disease^71^, where hippocampal hyperexcitability^72^ and altered network connectivity^73^ in patients with risk alleles for Alzheimer’s disease precedes the development of neuronal loss and dementia^74,75^. Evidence for this is seen in the significant increase in the prevalence of seizures observed in early stage AD patients^76^, and these seizures can hasten cognitive decline^77^. A more complete understanding of the signaling pathways which govern E-I balance may form a basis for the development new therapeutic strategies for the treatment of epilepsy, Alzheimer’s disease or other conditions in which an E-I imbalance exists. While the contribution of GABA_B_ autoreceptors to E-I balance is clear, the studies contained in this manuscript provide new insight on the mechanisms through which these receptors act to reduce GABA release outside of any pathophysiological context.

## MATERIALS AND METHODS

### Viral transduction of mouse parenchyma

Young male C57BL6/J animals of at least 17 grams were prepared from heterozygote breedings and genotyped for *Snap25* via genomic PCR of tail snip DNA (Transnetyx). Mice underwent anesthesia with 2.5% isoflurane (Somnosuite Unit, Kent Scientific), and the dorsal scalp skin was prepared through depilation and subdermal administration of 0.125% bupivacaine before placement in a stereotaxic frame (Stoelting). Meloxicam (2 mg/kg, s.c.) and ampicillin (10 mg/kg, s.c.) served as analgesic and antibiotic agents, respectively. Vaporized isoflurane anesthesia was continued throughout the surgical procedure, and body temperature was maintained at ∼37°C with heating elements on the stereotaxic frame. Following a ∼1 cm midline scalp incision and drilling of a 1mm burr hole over the injection site, a 35G needle attached to a 10 μl Nanofil syringe, controlled by a UMPI perfusion pump (WPI) was lowered to the following coordinates (in mm, relative to bregma): -2.15 A/P, ±1.5 M/L and -13 D/V, positioned parallel to the lateral axis. Viral solution (500 nL) was delivered at 250 nL/min, followed by a 5-minute dwell time. The needle was slowly retracted at 1 mm/minute. Wound clips were used to close the incision. Animals recovered individually on a heating pad and were injected with 60 units of normal saline s.c. before being returned to their home cage. Staples were removed 10 days post-surgery. A minimum 6-week period was permitted for AAV expression before sacrifice and preparation of acute slices.

### Preparation of acute mouse hippocampal slices

SNAP25Δ3 homozygote or WT littermates were anaesthetized with isoflurane and perfused intracardially with a solution of 93 mM N-methyl-d-glucamine, 2.5 mM KCl, 1.2 mM NaH2PO4, 20 mM HEPES, 10 mM MgSO4, 0.5 mM CaCl2, 25 mM d-glucose, 5 mM ascorbate, and 3 mM pyruvate, bubbled with 95% O2 and 5% CO2, pH7.4. Hippocampi were then dissected and slices 200 μm thick were produced via Vibratome (Leica VT1200, Leica Microsystems), and the solution was exchanged for ACSF (124 mM NaCl, 26 mM NaHCO3, 1.25 mM NaH2PO4, 3 mM KCl, 2 mM CaCl2, 1 mM MgCl2, and 10 mM d-glucose, bubbled with 95% O2 and 5% CO2.), for 30 minutes at 34 C post-sacrifice, before being returned to 25 C. Slices were maintained for a maximum of six hours post-sacrifice.

#### Whole cell patch clamp electrophysiology

Recordings were performed using slices secured by a stainless steel harp in a recording chamber superfused with constant flow of ACSF and warmed to 30°C. Pyramidal CA1 neuron bodies were visually identified and voltage-clamped using glass pipettes (4-6MΩ) filled halfway with 1mM QX314 in cesium-based patch solution (in mM): 146 CsMeSO_3_, 5 EGTA, 5 HEPES, 1 CaCl_2_, 1 NaCl_2_, 0.5 MgCl_2_, pH adjusted to 7.2 with CsOH. Osmolarity was adjusted to 290mOsm with ddH_2_O or mannitol. Patched cells were held at -50.0mV. Slices were treated with 5µM CNQX and 50µM AP5 in recording solution to isolate IPSCs and block polysynaptic activation.

For miniature IPSCs, cells were patched with a high-chloride patch solution (in mM): 146 KCl, 5 EGTA, 5 HEPES, 1 CaCl2, 1 NaCl2, 0.5 MgCl2, pH adjusted to 7.2 with KOH. Cells were held at -70.0mV. 1µM TTX, 5µM CNQX, and 50µM AP5 were added to the recording solution to prevent action potentials and isolate GABAergic signals.

For evoked IPSCs stimuli were evoked using a twisted bipolar nichrome electrode over interneurons in the stratum radiatum within 200mm of the clamped cell at various frequencies controlled by a Master-8 stimulator (AMPI). Responses were captured and digitized using an Axopatch 200B (Axon Instruments) integrating patch clamp amplifier and recorded using AxoGraph acquisition software. Drug treatments were allowed 20mL of perfusion over the slice prior to recording. Baclofen and CGP 55845 were prepared as 10mM stocks in water and DMSO, respectively, and stored at -20 C.

#### Epifluorescence imaging of acute slices

Acute slices virally transduced with AAV-mDlx-jGCaMP8f or AAV-syn-iGABASnFR2-WPRE were secured via harp in a temperature-stabilized flow chamber (30°C) filled with ACSF containing 50mM AP5 and 5mM CNQX to prevent polysynaptic activity. Solutions were bubbled with O_2_/CO2. A twisted-pair stimulating electrode was positioned over the Schaffer collaterals and depolarizing stimuli of 200-800 mA were administered for a duration of 50ms. Fields containing CA1 stratum pyramidale and stratum radiatum were imaged at a rate of 100 Hz. jGCaMP8f was excited via a 473nm LED and fast rises in 535nm fluorescence were captured on a 40x water-immersion objective using a pco.edge sCMOS camera with a Chroma 490-575 610-730 dual-bandpass filter affixed to an Olympus BX50WI fluorescence microscope. Micro-Manager 2^78^ was used for control of microscope hardware and imaging.

#### Lattice light sheet imaging of acute slices

Lattice light sheet imaging was performed according to previously published methods^45^ on a custom microscope based on an existing design from the laboratory of Eric Betzig^31^ (Janelia Research Campus). Virally transduced slices were prepared and via an identical manner to the preceding paragraph, secured to a raised platform positioned between excitation and recording objectives on a custom poly(methyl methacrylate) stage warmed to 30°C. GABAergic axonal structures in CA1 stratum radiatum and stratum pyramidale were visualized at 535nm subsequent to excitation with a 488nm laser, within single planes 512x512 pixels in size, recorded at 61.8 Hz. Highly fluorescent structures adjacent to the electrode with morphology resembling presynaptic terminals were located and stimuli were administered. Puncta within axonal structures that exhibited rapid increases in fluorescence subsequent to the application of stimuli were analyzed using ImageJ, and time-dependent dF/F ratios were obtained.

### Statistics

Electrophysiological data was acquired and exported using AxoGraph. Imaging data was analyzed using Fiji^79^. Wavemetrics Igor Pro and GraphPad Prism were utilized to fit exponential decay curves, conduct statistical tests, and plot data points.

## ACKNOWLEDGEMENTS

AAV1-syn-iGABASnFr2-WPRE was a generous gift from the GENIE project at Janelia Research Campus and the laboratories of Loren Looger and Jonathan Marvin. Other AAV constructs and preparations were made by VectorBuilder, Inc. We express gratitude to the research support team at the University of Illinois Chicago, including Ashley Miller, Lorissa Lamoureux, DVM, Jason York, Ph.D, and Justin Jania, JD.

