## Supplemental Text and Figures for "Presynaptic GABA_B_ autoreceptors suppress neurotransmitter release during repetitive stimulation via the Gβγ-SNARE pathway"

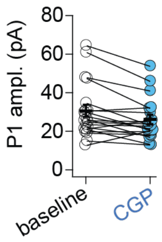


**Supplemental Figure 1. $GABA_B$ antagonists do not alter IPSC P1 amplitudes.** Scatter plot of the first IPSC amplitude in the train before and after CGP 55845. The difference was not significant, tabulated via unpaired two-tailed Student's t-test. Error bars represent means + S.E.M.

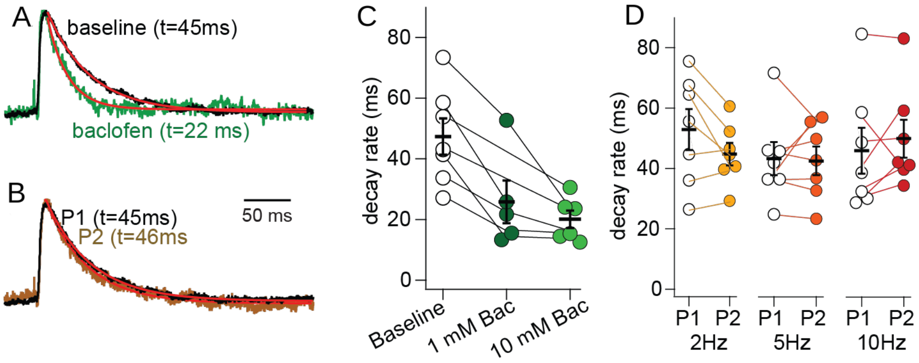


**Supplemental Figure 2. Baclofen inhibits IPSC decay constants.** A, B: Representative trace of IPSC decay rates in the presence or absence of baclofen (green). Time intervals utilized to calculate τ are highlighted in red. CScatterplot of decay constants in the presence or absence of baclofen. Representative traces of IPSC decay rates for the first (black) or the final IPSC (gold) in the train. D,. Scatterplot of decay constants for the first and second IPSC at 2, 5, or 10 Hz. P-values shown tabulated by paired two-tailed Student's t-test. Error bars represent means + S.E.M.

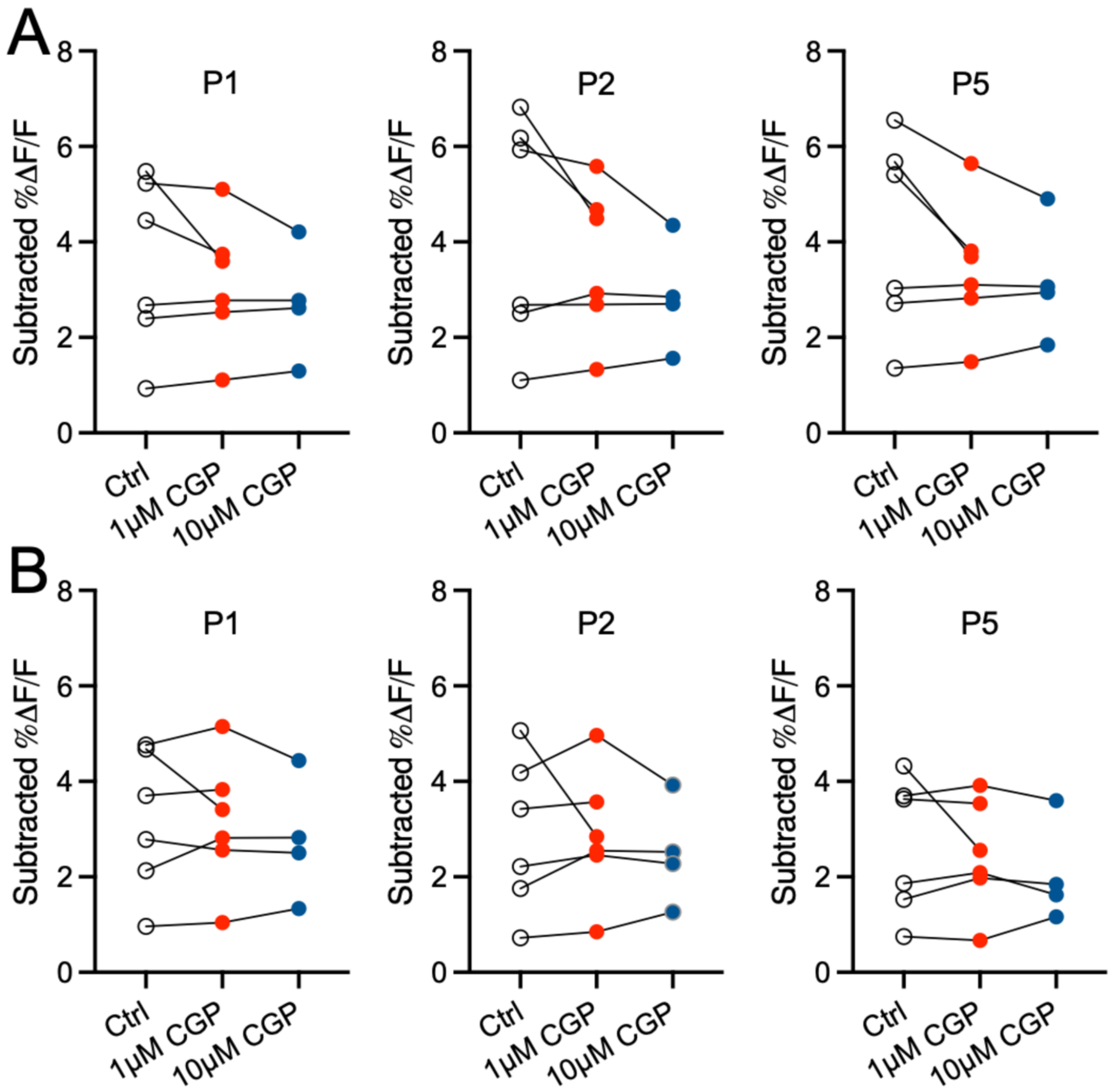


**Supplemental Figure 3: GABAB antagonists do not alter the height of any mDlx-jGCaMP8f flux in trains of stimuli.** A. Scatter plot of subtracted peak amplitudes for the first, second or fifth stimulus in the train at 2Hz, in the presence or absence of 1 or 10 μM CGP 55845. B. Scatter plot of subtracted peak amplitudes as above, at 10Hz. P-values shown tabulated by paired two-tailed Student's t-test.

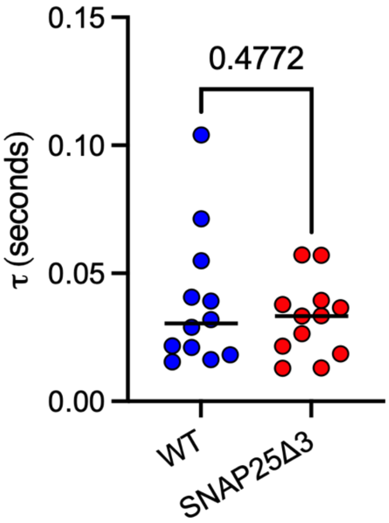


**Supplemental Figure 4: The SNAP25D3 mutation does not affect IPSC decay rates.** Scatterplot of decay constants in WT and SNAP25D3 for the decay rate of the fifth and final stimuli in the train at 2 Hz, as in Supplemental Figure 2. P-values shown tabulated by unpaired two-tailed Student's t-test. Error bars represent means + S.E.M.

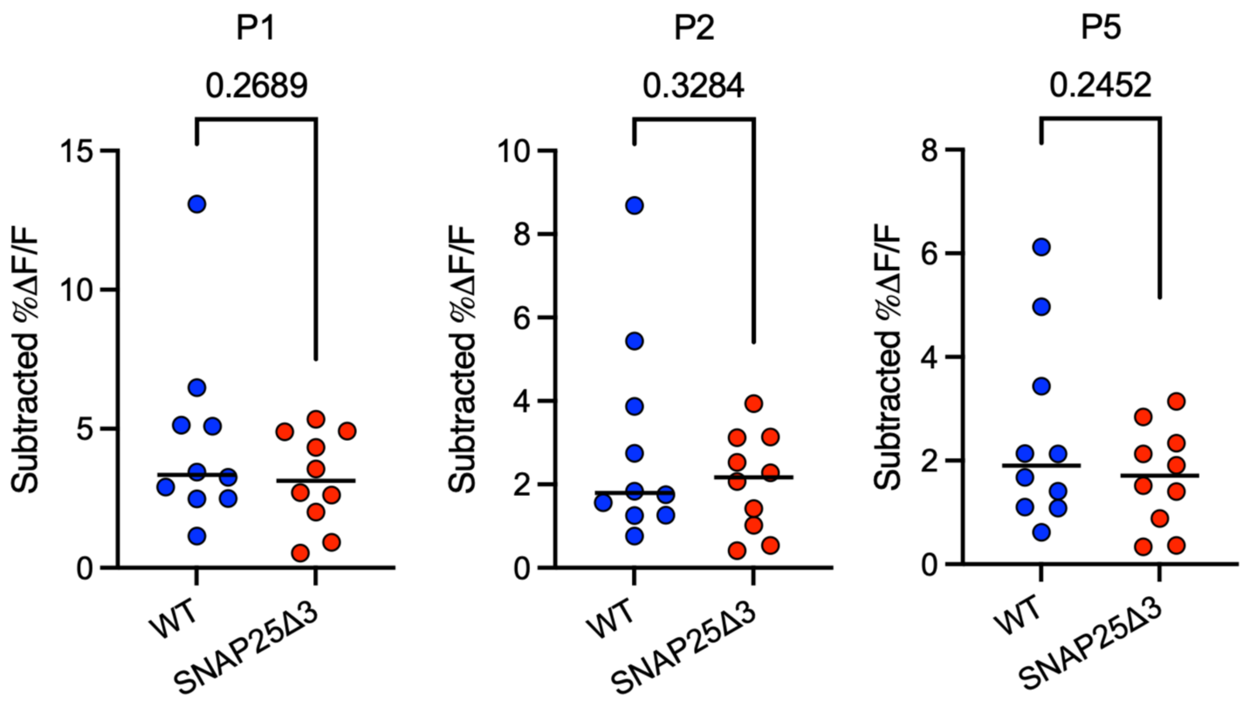


**Supplemental Figure 5: The effect of genotype on individual iGABASnFr2 peak heights.** Scatter plot of subtracted peak amplitudes for evoked WT and SNAP25D3 iGABASnFr2 for the first, second or fifth stimulus in the train at 2Hz. P-values shown tabulated by unpaired two-tailed Student's t-test. Error bars represent means + S.E.M.

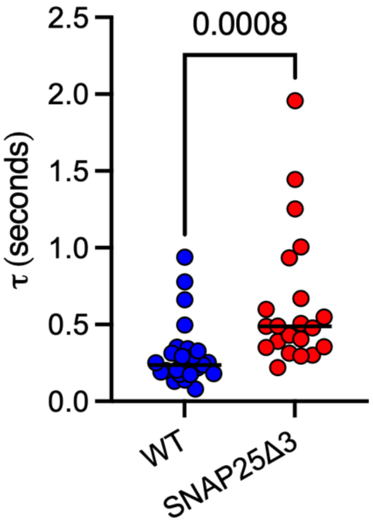


**Supplemental Figure 6: iGABASnFr2 decay rates are much slower in SNAP25Δ3.** Scatterplots comparing the effect of SNAP25Δ3 genotype on the decay constant for iGABASnFr2 fluorescence after the fifth stimuli in the train at 2 Hz. Displayed P-values tabulated via unpaired two-tailed Student's t-test. Error bars represent means + S.E.M.

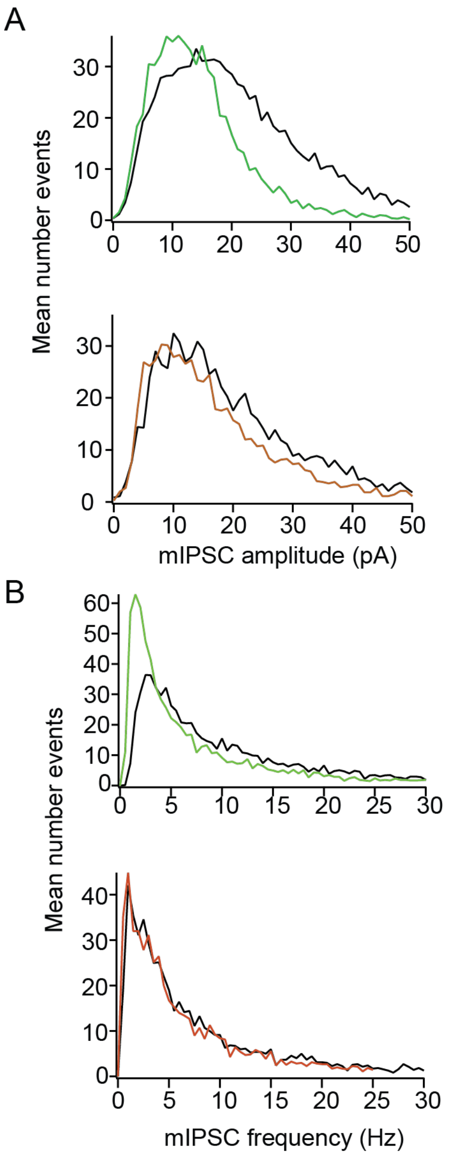


**Supplemental Figure 7: Baclofen produces a leftward shift in the distribution of miPSC amplitudes and frequencies in WT animals, but to a lesser extent in SNAP25Δ3 animals.** A. Histogram of individual mIPSC amplitudes from WT neurons or SNAP25Δ3 neurons in the presence or absence of baclofen (green and orange for WT and SNAP25Δ3, respectively). Baclofen produces a leftward shift in the distribution of miPSC amplitudes and frequencies. B. Histogram of individual mIPSC frequencies from SNAP25Δ3 neurons in the presence or absence of baclofen. Effect of baclofen on the distribution of amplitudes and frequencies was tabulated by the two-sample Kolmogorov-Smirnov test.
